# Identification of inosine monophosphate dehydrogenase as a potential target for anti-monkeypox virus agents

**DOI:** 10.1101/2022.12.26.521968

**Authors:** Takayuki Hishiki, Takeshi Morita, Daisuke Akazawa, Hirofumi Ohashi, Eun-Sil Park, Michiyo Kataoka, Junki Mifune, Kaho Shionoya, Kana Tsuchimoto, Shinjiro Ojima, Aa Haeruman Azam, Shogo Nakajima, Tomoki Yoshikawa, Masayuki Shimojima, Kotaro Kiga, Ken Maeda, Tadaki Suzuki, Hideki Ebihara, Yoshimasa Takahashi, Koichi Watashi

**Author notes:** These authors contributed equally to this work (Takayuki Hishiki, Takeshi Morita). These authors also contributed equally to this work (Daisuke Akazawa, Hirofumi Ohashi). Corresponding author: Koichi Watashi, Ph.D.

## Abstract

Monkeypox virus (MPXV) is a neglected zoonotic pathogen that caused a worldwide outbreak in May 2022. Given the lack of an established therapy, the development of an anti-MPXV strategy is of vital importance. To identify drug targets for the development of anti-MPXV agents, we screened a chemical library using an MPXV infection cell assay and found that gemcitabine, trifluridine, and mycophenolic acid (MPA) inhibited MPXV propagation. These compounds showed broad-spectrum anti-orthopoxvirus activities and presented lower 90% inhibitory concentrations (0.032-1.40 μM) than brincidofovir, an approved anti-smallpox agent. These three compounds have been suggested to target the post-entry step to reduce the intracellular production of virions. Knockdown of inosine monophosphate dehydrogenase (IMPDH), the rate-limiting enzyme of guanosine biosynthesis and a target of MPA, dramatically reduced MPXV DNA production. Moreover, supplementation with guanosine recovered the anti-MPXV effect of MPA, suggesting that IMPDH and its guanosine biosynthetic pathway regulate MPXV replication. By targeting IMPDH, we identified a series of compounds with stronger anti-MPXV activity than MPA. These evidences propose that IMPDH is a potential target for the development of anti-MPXV agents.

**Importance:** Monkeypox is a zoonotic disease caused by infection with the monkeypox virus, and a worldwide outbreak occurred in May 2022. The smallpox vaccine has recently been approved for clinical use against monkeypox in the United States. Although brincidofovir and tecovirimat are drugs approved for the treatment of smallpox by the U.S. Food and Drug Administration, their efficacy against monkeypox has not been established. Moreover, these drugs may present negative side effects. Therefore, new anti-monkeypox virus agents are needed. This study revealed that gemcitabine, trifluridine, and mycophenolic acid inhibited monkeypox virus propagation, exhibited broad-spectrum anti-orthopoxvirus activities. We also suggested inosine monophosphate dehydrogenase as a potential target for the development of anti-monkeypox virus agents. By targeting this molecule, we identified a series of compounds with stronger anti-monkeypox virus activity than mycophenolic acid.

## Introduction

Monkeypox is a zoonotic disease caused by infection with the monkeypox virus (MPXV). MPXV is an enveloped virus with a double-stranded DNA genome of approximately 190 kb in length. It belongs to the genus *Orthopoxvirus* of the family *Poxviridae*, which includes the smallpox, vaccinia, and cowpox viruses and other animal-associated poxviruses (1). The natural hosts of MPXV are most likely rodents, and MPXV is transmitted to humans by infected animals through bites or contact with the blood or body fluids. MPXV is also transmitted through human-to-human contact via droplets or body fluids (39). Starting in May 2022, wide-scale monkeypox cases were reported in multiple countries where this disease had not been previously endemic, and by November 2022, more than 80,000 cases of infection had been reported in over 110 countries, mainly in Europe and the United States, with most of these infections transmitted via sexual contact (40). Given the current status of MPXV and possible future outbreaks, medical countermeasures and further research on MPXV should be developed.

The smallpox vaccine has recently been approved for clinical use against monkeypox in the United States (41). Brincidofovir and tecovirimat are drugs approved for the treatment of smallpox by the U.S. Food and Drug Administration (FDA) under the agency’s animal rule (2). Brincidofovir is a lipid conjugate of cidofovir, a nucleoside analog active against cytomegalovirus, and it suppresses viral genome replication by inhibiting viral DNA polymerase (3–7). The efficacy of brincidofovir on monkeypox has not been established, and a recent clinical report showed no clinical benefit to monkeypox patients and rather indicated liver toxicity by brincidofovir (2, 8). Tecovirimat is an FDA-approved anti-smallpox drug that inhibits virion maturation; however, its clinical efficacy against monkeypox is poorly documented because of the limited chance of clinical treatment (2, 8–10, 38). In cell culture studies, tecovirimat treatment has been reported to induce drug-resistant viruses (11, 12), although the clinical drug resistance profile is not clear. Thus, development of new anti-MPXV strategy would provide alternative therapeutic options.

In this study, we aimed to identify a new drug target for MPXV. We screened a chemical compound library using an MPXV infection cell culture assay and found that gemcitabine, trifluridine, and mycophenolic acid (MPA) inhibited MPXV replication. An analysis of the anti-MPXV activity of MPA showed that inosine monophosphate dehydrogenase (IMPDH) and the guanine nucleotide biosynthesis pathway have significant roles in regulating MPXV replication. By targeting IMPDH, we identified compounds with more potent anti-MPXV activity than MPA. Therefore, we propose IMPDH as a potential target for the development of anti-MPXV agents.

## Results

### Anti-MPXV activity of gemcitabine, trifluridine, and mycophenolic acid

To identify compounds that inhibit MPXV propagation, we screened 121 compounds previously reported to have anti-vaccinia virus activity; however, most of their modes of action are unknown (13) (Table S1). On the first screen, VeroE6 cells were infected with MPXV Zr-599 (Congo Basin strain) at a multiplicity of infection (MOI) of 0.1 for 72 h in the presence of 10 μM of each compound, with the exception of two compounds treated at 2 μM (Table S1). The cytopathic effect induced by MPXV propagation was detected by observing cell morphology using a microscope (Fig. 1A) and quantifying cell viability using a high-content imaging analyzer (Fig. S1) (14). Although cells remained confluent without virus inoculation, inoculation with MPXV induced extensive cell death after 72 h (Fig. 1A-a, b). As positive controls, treatment with tecovirimat and brincidofovir protected cells from MPXV-induced cytopathic effects and augmented the number of surviving cells to 223 and 103 fold, respectively (Fig. 1A-c, d, and S1). The screening revealed 74 compounds that increased the survival cell number by more than 50 fold relative to that of the DMSO-treated control cells (Fig. S1).

**Fig. 1.**
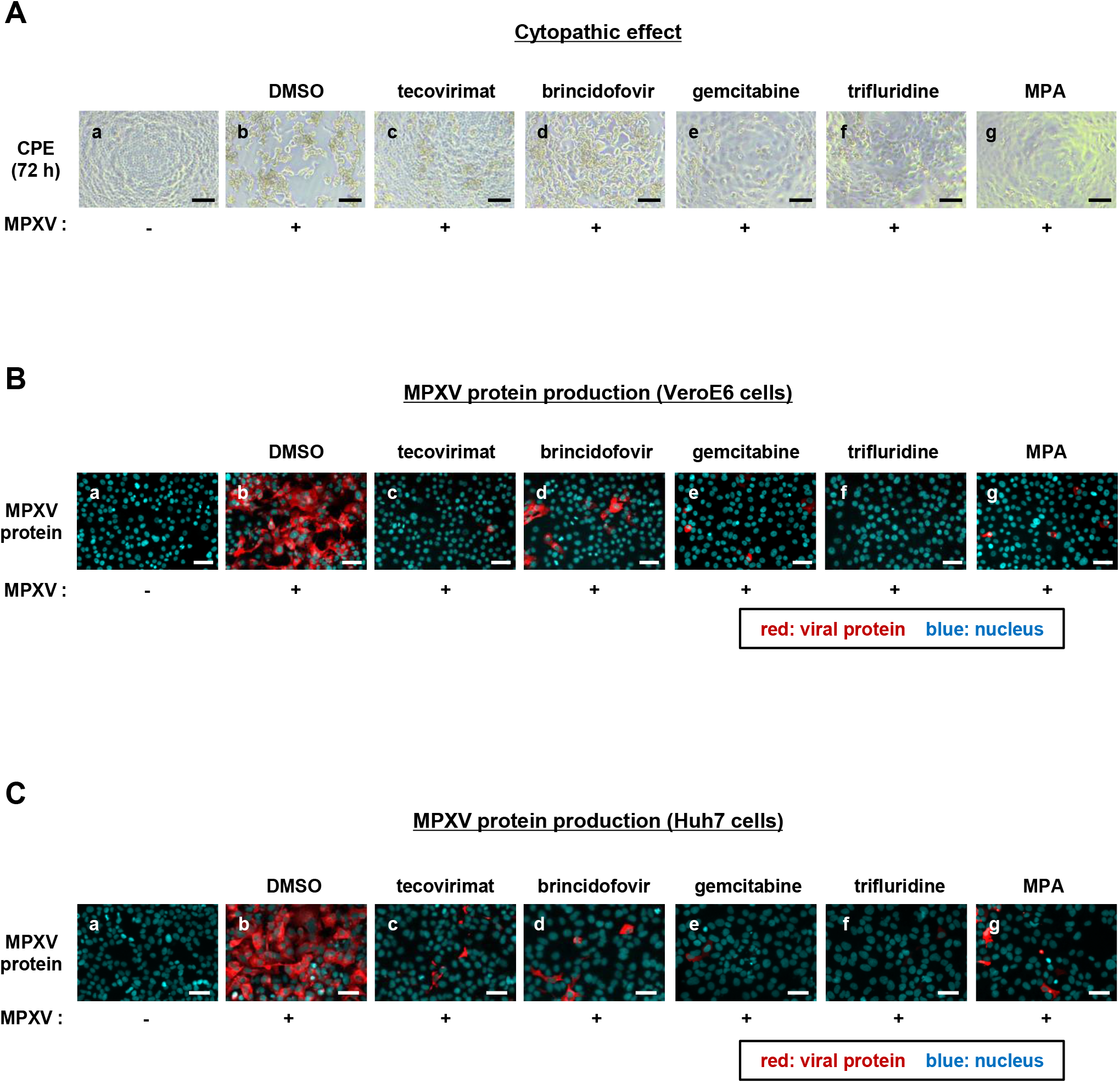
Anti-MPXV activity of mycophenolic acid, gemcitabine, and trifluridine. (A) VeroE6 cells were infected with (b-g) or without (a) MPXV at an MOI of 0.1 and then treated with the indicated compounds at 10 μM or DMSO at 0.1% (b). The panels show the cell morphology at 72 h post-infection via microscopy. Scale bars, 100 μm. (B) VeroE6 cells were infected with MPXV at an MOI of 0.1 in the presence of either 0.1% DMSO, 40 nM tecovirimat, 5 μM brincidofovir, 0.1 μM gemcitabine, 5 μM trifluridine, or 5 μM mycophenolic acid (MPA). At 24 h post-infection, cells were harvested for immunfluorescence analysis to detect viral proteins and nuclei. Red, MPXV protein; blue, nuclei. Scale bars, 50 μm. (C) Huh7 cells were infected with the same amount of MPXV as inoculum as shown in Fig. 1B in the presence of 5 μM of the indicated compounds or 0.1% DMSO. At 24 h post-infection, the cells were harvested for immunfluorescence analysis as shown in Fig. 1C. Scale bars, 50 μm.

Among the hit compounds, we focused on the three compounds gemcitabine, trifluridine, and mycophenolic acid (MPA) (Fig. 1A-e, f, g) because they have been reported to inhibit the replication of multiple virus species, such as adenovirus, herpes simplex virus, Zika virus, severe acute respiratory syndrome coronavirus 2 (SARS-CoV-2), and dengue virus (15–19). We confirmed the anti-MPXV activity of the three compounds by detecting viral protein production in MPXV-infected cells by immunofluorescence analysis. VeroE6 cells infected with MPXV at an MOI of 0.1 for 1 h were incubated with each compound for another 23 h and then fixed to detect anti-MPXV protein together with DAPI for nuclear staining. The cells did not show cytopathology under these conditions at 24 h after virus inoculation (Fig. 1B, blue). As shown in Fig. 1B, MPXV protein expression was drastically reduced upon treatment with gemcitabine, trifluridine, and MPA (Fig. 1B-e, f, and g), which was similar to results obtained for tecovirimat and brincidofovir (Fig. 1B-c and d). These antiviral effects were also observed in the human-derived cell line Huh7 cells (Fig. 1C), suggesting that the anti-MPXV activity of these compounds was not dependent on the cell type.

To examine the activity of these compounds against multiple orthopoxviruses, we analyzed their effect on infection assays using the MPXV Liberia strain (West African strain), vaccinia virus, and cowpox virus. As shown in Fig. 2, gemcitabine, trifluridine, and MPA clearly reduced the expression of viral proteins in cells inoculated with all viruses (Fig. 2). These results suggest that gemcitabine, trifluridine, and MPA possess antiviral activities against a wide range of orthopoxviruses.

**Fig. 2.**
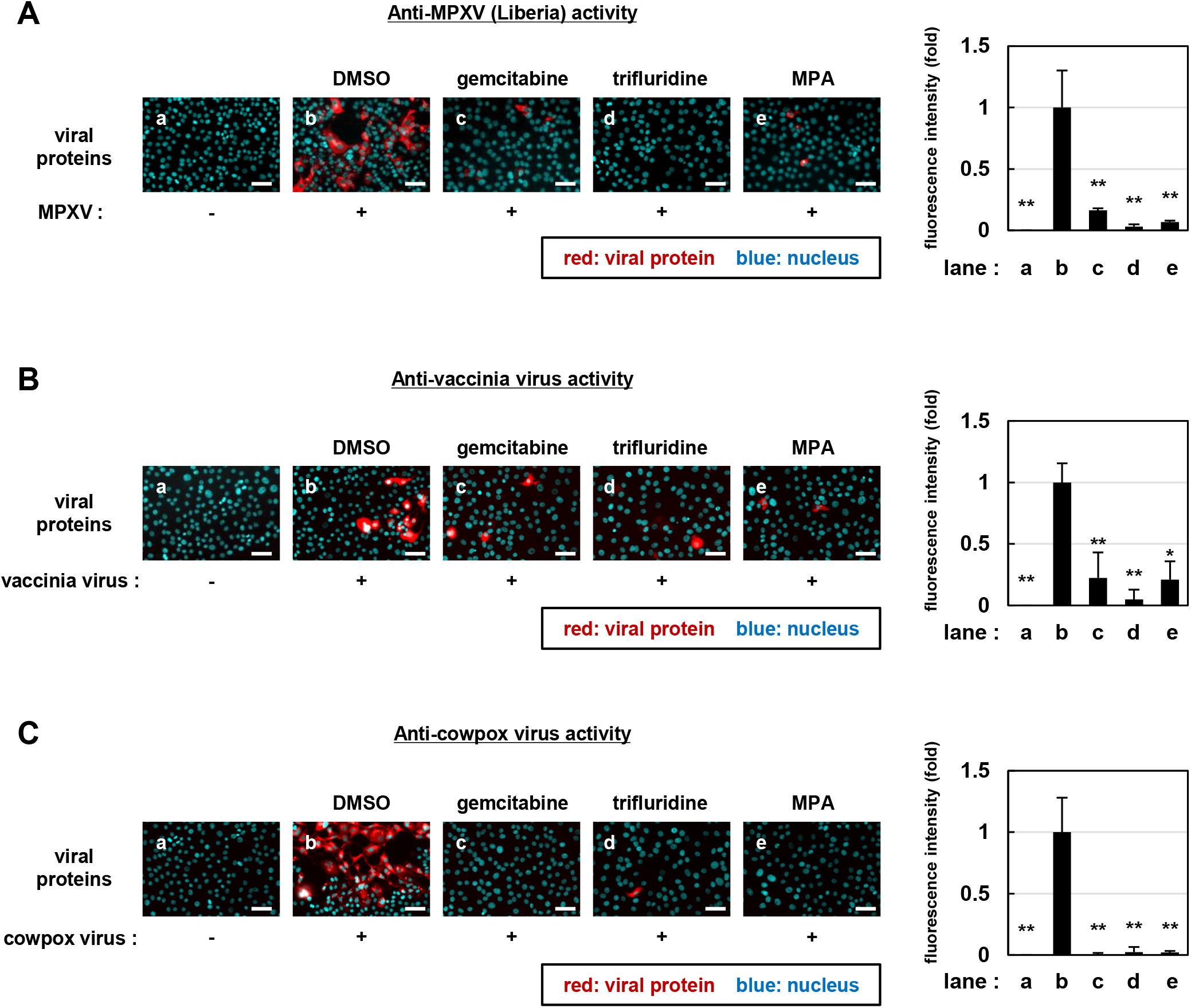
Broad spectrum anti-orthopoxvirus activity of gemcitabine, trifluridine, and MPA. VeroE6 cells infected with the MPXV Liberia strain (A), vaccinia virus (B), and cowpoxvirus (C) at an MOI of 0.1 and co-cultured with 5 μM of each compound. At 24 h post-infection, cells were harvested for immunfluorescence analysis to detect viral proteins and nuclei. Red, MPXV protein; blue, nuclei. Scale bars, 50 μm. Quantification of red fluorescence area calculated by Dynamic Cell Count (Keyence) and shown relative to that of the DMSO-treated control cells (right graph).

### Dose response curve of anti-MPXV activity for gemcitabine, trifluridine, and MPA

To quantify the anti-MPXV activity of the three compounds, VeroE6 cells infected with MPXV (MOI of 0.03) for 1 h were incubated with varying concentrations of the compounds up to 10 μM for an additional 29 h to assess intracellular viral DNA levels by real-time PCR and examine cytotoxicity by water-soluble tetrazolium salt (WST) assay. Brincidofovir was analyzed under the same conditions as the positive control (20). All compounds reduced the levels of MPXV DNA in a dose-dependent manner (Fig. 3B) and did not show significant cytotoxicity (Fig. 3C). The 50% and 90% maximal inhibitory concentrations (IC_50_ and IC_90_) as well as the 50% maximal cytotoxic concentrations (CC50) of each compound are shown in Fig. 3B and C. These results indicated that the three compounds exhibited dose-dependent anti-MPXV activity without cytotoxicity and presented a lower IC_90_ than brincidofovir.

**Fig. 3.**
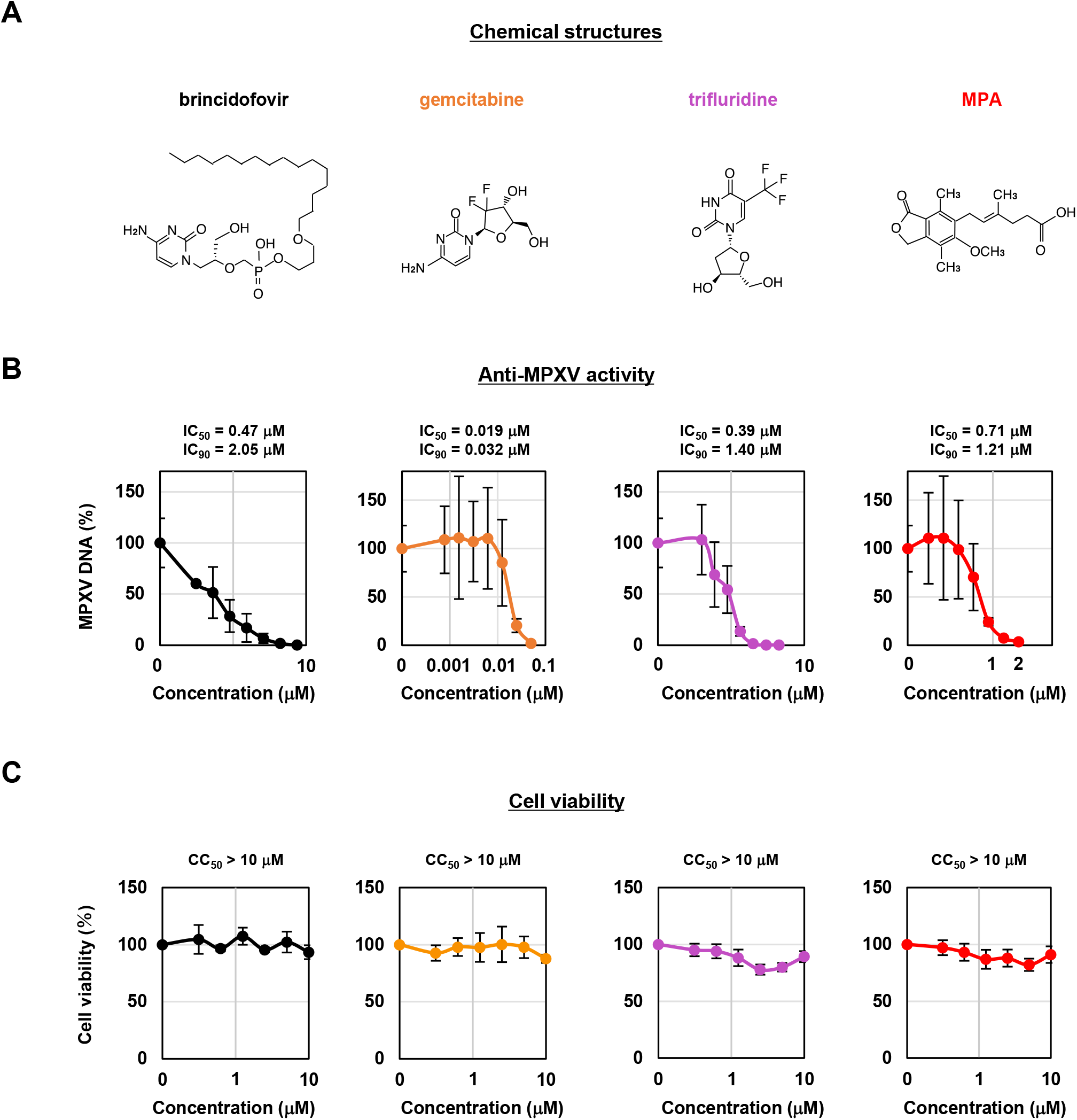
Dose-response curve for anti-MPXV activity of gemcitabine, trifluridine, and MPA. (A) Chemical structure of brincidofovir, gemcitabine, trifluridine, and MPA. (B) VeroE6 cells were infected with MPXV at an MOI of 0.03 for 1 hour and then incubated with the media supplemented with the indicated concentrations of compounds for 30 h. Intracellular viral DNA was measured by real-time PCR. The y-axis shows the value relative to that for DMSO-treated cells as a control. IC_50_ and IC_90_ calculated by a linear regression are presented above the graphs. (C) VeroE6 cells incubated with the indicated concentrations of compound for 30 h were subjected to the detection of cell viability. The y-axis shows values relative to that of the DMSO-treated cells as a control. CC50 values calculated by linear regression are also represented.

### Gemcitabine, trifluridine, and MPA target the post-entry phase in MPXV life cycle

A schematic diagram of the MPXV life cycle is shown in Fig. 4A. MPXV attaches to the surface of a target cell to enter intracellularly, where the viral core is delivered into the cytoplasm (entry phase, Fig. 4A). Through early transcription, protein synthesis, and uncoating of the core, viral DNA replicates and drives intermediate and late transcription, which is followed by viral assembly in a specific compartment called the cytoplasmic viral factory, and further stepwise virion maturation produces infectious virions (post-entry phase, Fig. 4A) (21). To clarify the phase of the MPXV life cycle that is inhibited by the compounds, we performed a time-of-drug-addition assay, in which the entry and post-entry phases are distinguished by changing the treatment time of the compound (Fig. 4B, left) (14, 22). Compounds were added at different times: over the entire assay period of 24 h (a: whole life cycle), within the initial 2 h (b: entry phase), or over the last 22 h after viral infection (c: post-entry and re-infection phase) (Fig. 4B, left). We assessed the antiviral activity by detecting intracellular viral DNA using real-time PCR for each condition. Brincidofovir, a positive control that inhibits viral genome replication, showed significant antiviral activity in conditions (a) and (c) but not in condition (b) (Fig. 4B, right) (20). In contrast, heparin, which is reported to inhibit viral entry, showed significant antiviral activity under condition (b) (Fig. 4B, left). These results indicate that our time-of-drug-addition assay could successfully distinguish entry inhibitors from those that inhibit viral replication. As shown in Fig. 4B, gemcitabine, trifluridine, and MPA were significantly reduced under conditions (a) and (c) but not under condition (b), suggesting that these compounds inhibit the post-entry phase in the MPXV life cycle.

**Fig. 4.**
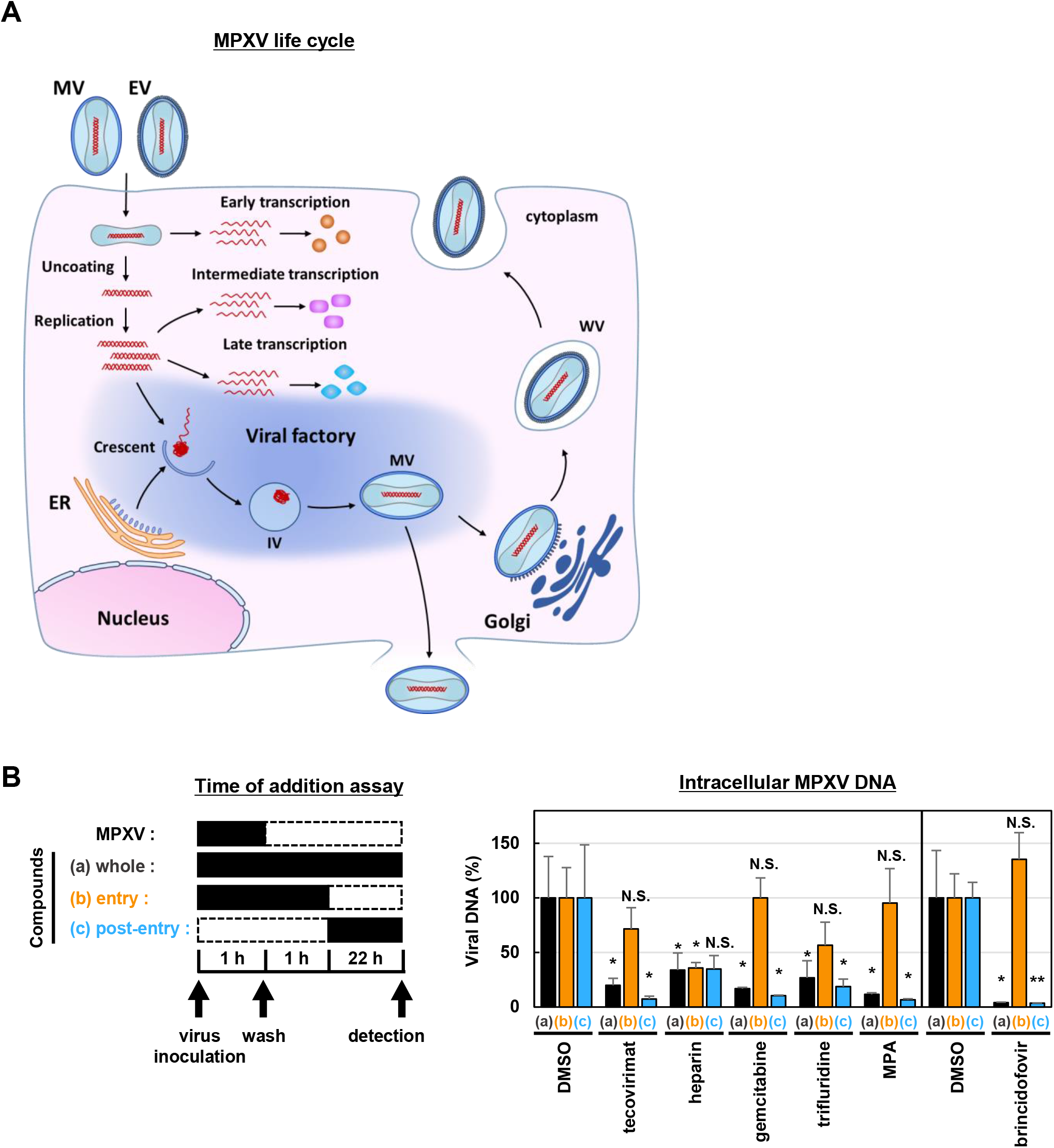
Gemcitabine, trifluridine, and MPA inhibit viral post-entry phase. (A) Schematic representation of the MPXV life cycle. (B) Left, schematic representation of the time-of-drug-addition assay. VeroE6 cells were infected with MPXV at an MOI of 0.03 for 1 hour with (a, b) or without (c) the compound. After washing out the inoculated MPXV, the cells were incubated with the media with (a, b) or without (c) the compound for 1 hour. After washing again, the cells were further incubated with the media with (a, c) or without (b) the compound for 22 h. In summary, the cells were treated with the compound for 0–24 (a: whole life cycle), 0–2 (b: entry phase), or 2–24 h (c: post-entry and re-infection phase) post-MPXV infection. Black and dotted boxes indicate the periods of treatment and non-treatment, respectively. Right: anti-MPXV activity was examined by quantifying viral DNA in cells by real-time PCR. The y-axis shows the value relative to that of the DMSO-treated cells. Statistical significance against DMSO treated cells is shown (\**P*; < 0.05, \*\**P*; < 0.01, N.S.; not significant).

### Observation of intracellular structures by electron microscopic analysis

Poxvirus infection induces the formation of an intracellular structure called the cytoplasmic factory, which represents a hallmark of infected cells and is involved in virion assembly (23, 24). We then observed the intracellular morphological features of the compound-treated cells by transmission electron microscopy. Tecovirimat, a particle maturation inhibitor, was used as the positive control (25). Crescents, immature, mature, and wrapped particles were observed in DMSO-treated MPXV-infected cells (Fig. 5A-b, c), while crescents, immature, and mature virions but not wrapped virions were observed in tecovirimat-treated cells (Fig. 5B-d, h), which is consistent with the mode of action of tecovirimat. In contrast, few virions were observed in gemcitabine-, trifluridine-, and MPA-treated cells (Fig. 5B-e, f, g, i, j, and k). These observations are consistent with the results obtained for gemcitabine, trifluridine, and MPA, which suppress the phase before virion assembly.

**Fig. 5.**
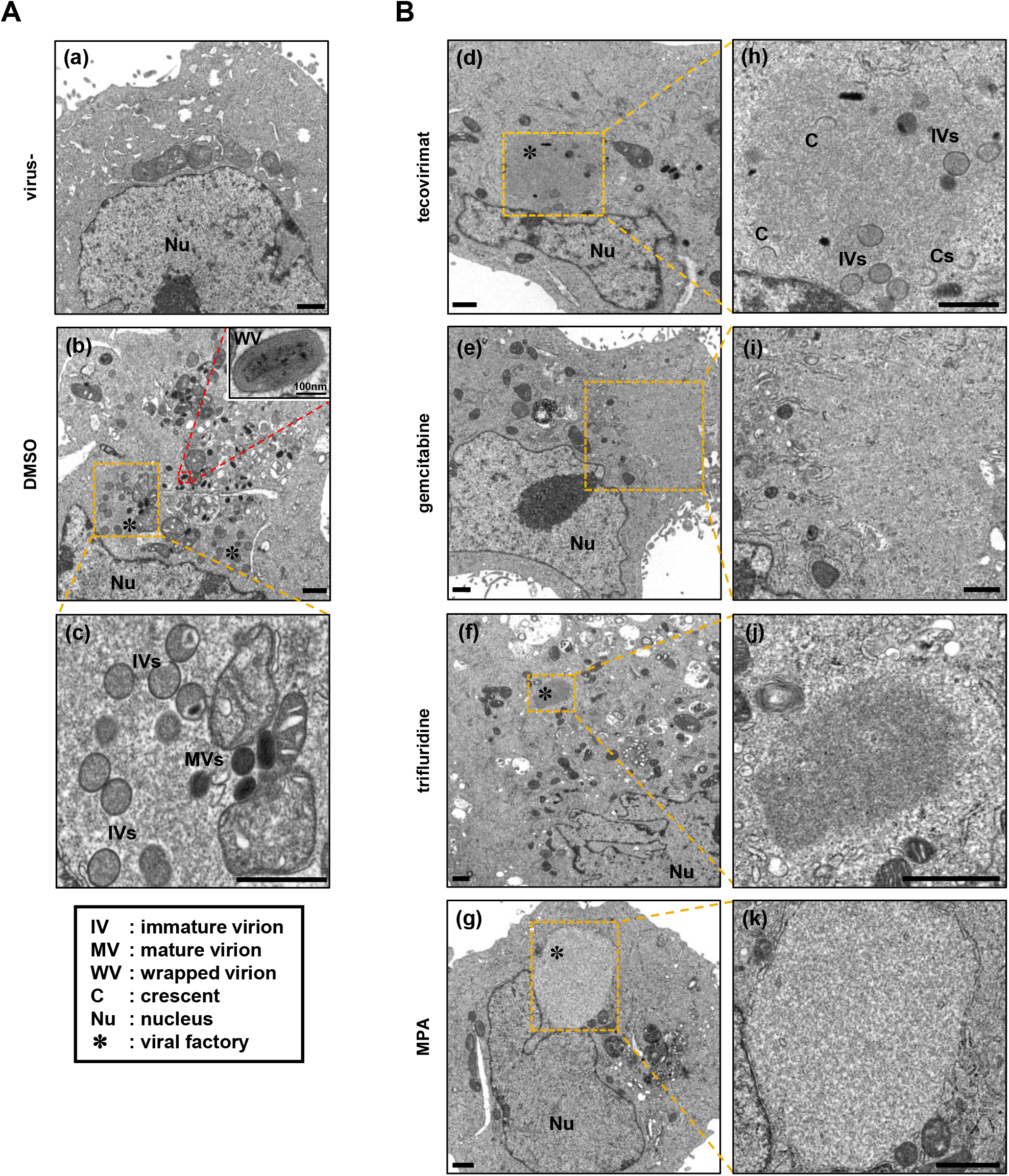
Electron microscopy observation of intracellular structures in compound-treated cells. (A, B) VeroE6 cells were infected (b to k) or uninfected (a) with MPXV and were treated with the indicated compounds (b, 0.1% DMSO; d, 5 μM tecovirimat; e, 5 μM gemcitabine; f, 5 μM trifluridine; g, 5 μM MPA). At 24 h post-treatment, the cells were fixed and processed for ultrastructural analysis by transmission electron microscopy as shown in the Methods. A total of 150 cells were observed for each sample, and representative images of the morphology are shown. The images in c, h, i, j, and k show-high magnification images of the yellow insets in b, d, e, f, and g, respectively. The inset in panel b shows a high-magnification image of the red frame area. Scale bars in a to k, 1 μm; scale bar for the upper right inset of panel b, 100 nm. IV, immature virion; MV, mature virion; WV, wrapped virion; C, crescent; Nu, nucleus; *, cytoplasmic viral factory.

### Mycophenolic acid suppresses MPXV replication through inhibition of IMPDH

Among the three identified compounds, gemcitabine and trifluridine are nucleoside analogs that are likely to target viral polymerase similar to brincidofovir (16–18, 20). Therefore, we analyzed the mechanism of action of MPA against MPXV MPA inhibits inosine monophosphate dehydrogenase (IMPDH), which is the rate-limiting enzyme of guanosine triphosphate (GTP) *de novo* synthesis (15, 19, 26) (Fig. 6A). IMPDH is composed of two isoforms: IMPDH1 and IMPDH2. To examine the role of IMPDH in MPXV replication, we transfected Huh7 cells with or without small interfering RNA (siRNA) targeting IMPDH1/2 or randomized control siRNA. At 48 h post-transfection, we confirmed the knockdown of both endogenous IMPDH1 and IMPDH2 at both the mRNA and protein levels (Fig. 6B-i, ii). At 48 h post-transfection with siRNA, we infected the cells with MPXV for 24 h and quantified the intracellular MPXV DNA by real-time PCR to examine the MPXV replication levels. As shown in Fig. 6B-iii, the MPXV DNA levels were significantly reduced in IMPDH1/2-depleted cells (Fig. 6B-iii).

**Fig. 6.**
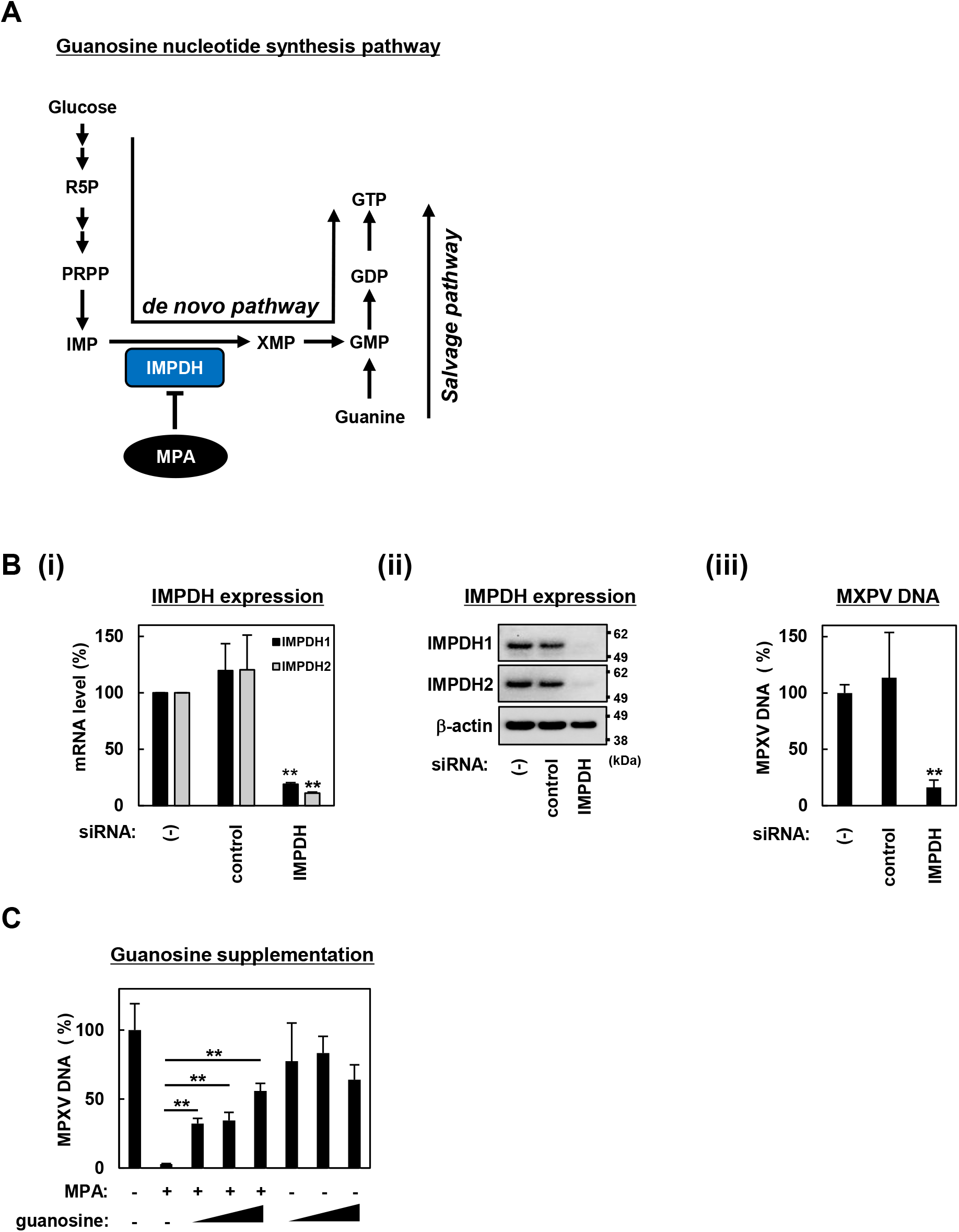
Essential role of IMPDH in the anti-MPXV activity of MPA. (A) Schematic representation of the guanosine nucleotide synthesis pathway. In the *de novo* pathway, an initial substrate, i.e., glucose, is converted to ribose-5-phosphate (R5P), phosphoribosyl diphosphate (PRPP), and then to inosine monophosphate (IMP) in a stepwise manner. IMPDH catalyzes the conversion from IMP to xanthine monophosphate (XMP) as a rate-limiting step of the pathway. XMP is converted through guanosine monophosphate (GMP) and guanosine diphosphate (GDP) to guanosine triphosphate (GTP). In the salvage pathway, GMP is also produced from guanine. (B) Huh7 cells were transfected with or without [(-)] siRNA targeting IMPDH (IMPDH) or randomized control siRNA (control). At 48 h post-transfection, intracellular RNA for IMPDH1 and IMPDH2 and the protein expression for IMPDH1, IMPDH2, and ß-actin were detected by real-time RT-PCR (i) and immunoblot analysis (ii), respectively. The y-axis in the Fig. 6B-i shows the value realtive to that for the untransfected cells. Upper, middle, and lower panels in Fig. 6B-ii show the protein production for IMPDH1, IMPDH2, and ß-actin, respectively. The positions for the molecular weight markers (62, 49, and 38 kDa) are also shown. Intracellular MPXV DNA levels at 72 h post-transfection with siRNA were quantified by real-time PCR and are shown as the percentage relative to that of the untransfected cells (iii). (C) Huh7 cells infected with the same amount of MPXV as inoculum in Fig. 1B were treated with or without 5 μM MPA and supplemented with or without varying amount of guanosine (12.5, 25, and 50 μM). After 24 h of treatment, intracellular viral DNA was measured by real-time PCR and is shown as the value relative to that for the DMSO-treated cells. Statistical significance is shown.

To further examine the relevance of the guanosine nucleotide synthetic pathway in the anti-MPXV activity of MPA, we performed a rescue experiment by complementing the MPA treatment with guanosine, a downstream product of the IMPDH-catalyzing step. MPXV-infected Huh7 cells were treated with MPA in the presence or absence of varying concentrations of guanosine and examined to detect viral DNA in cells at 24 h post-treatment. As shown in Fig. 6C, supplementation with guanosine clearly recovered the MPA-mediated reduction of viral DNA levels in a dose-dependent manner, suggesting that IMPDH and its guanosine synthesis pathway are targets for the observed anti-MPXV activity of MPA. Thus, IMPDH may be crucial for the efficient replication of MPXV.

### IMPDH inhibitors reduce MPXV propagation

The above results indicate that IMPDH is a potential target for the development of anti-MPXV agents. Therefore, the effects of known IMPDH inhibitors mycophenolate mofetil, AVN-944, merimepodib, and ribavirin were investigated. MPXV-infected Huh7 cells were incubated with these IMPDH inhibitors for 24 h to assess MPXV replication by detecting intracellular MPXV DNA levels by real-time PCR and cytotoxicity by the WST assay. All these IMPDH inhibitors clearly reduced MPXV DNA levels in a dose-dependent manner without showing cytotoxic effects (Fig. 7A and B), thus supporting the essential role of IMPDH in efficient MPXV replication. Based on the IC_50_ and IC_90_ values against MPXV shown in Fig. 7, mycophenolate mofetil, AVN-944, and merimepodib showed stronger anti-MPXV activity than MPA. Thus, targeting IMPDH would enable the identification of new anti-MPXV agents with high potency.

**Fig. 7.**
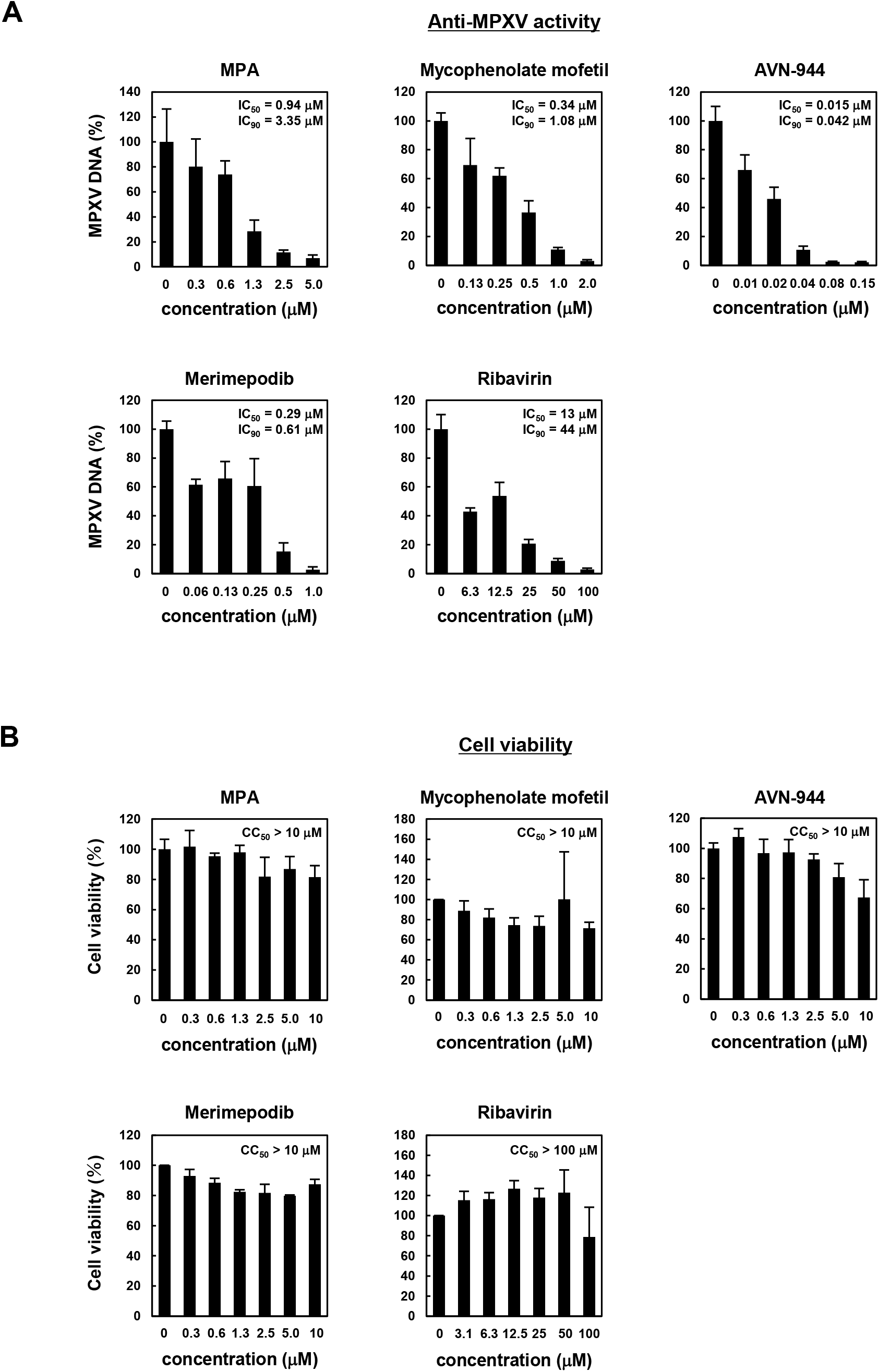
IMPDH inhibitors reduce the MPXV replication level. Huh7 cells infected with MPXV were treated with the indicated compounds and concentrations for 24 h. Intracellular MPXV DNA was measured by real-time PCR and the value relative to that of the DMSO-treated cells is shown. IC_50_ and IC_90_ were calculated by linear regression and are represented above the graphs. (B) Huh7 cells incubated with the indicated concentrations of compound for 24 h were subjected to a cytotoxicity assay to determine cell viability. The y-axis shows the values to that of DMSO-treated cells as a control.

## Discussion

In this study, we screened a compound library using an MPXV infection cell culture assay and identified 74 compounds as first hits. Among the hit compounds, we focused on the three compounds gemcitabine, trifluridine, and MPA and showed that they inhibited multiple strains of MPXV, vaccinia virus, and cowpox virus. Gemcitabine and trifluridine present anti-MPXV activities that are equivalent to or more potent than that of brincidofovir, and these nucleoside analogs are likely to show similar targeting of viral polymerization as brincidofovir and become incorporated into the viral genome or interfere with viral polymerase, resulting in the suppression of viral genome replication (18, 27, 28). Gemcitabine targets poliovirus RNA polymerase to inhibit viral replication (29). The ability of gemcitabine and trifluridine to target MPXV polymerization was supported by the fact that these compounds inhibited the post-entry phase and reduced intracellular virion accumulation.

MPA inhibits the enzymatic activity of IMPDH, the rate-limiting enzyme for the *de novo* synthesis of guanine nucleotides. In this study, we demonstrated that the anti-MPXV activity of MPA is mediated by the inhibition of IMPDH and the guanine synthetic pathway. Inhibition of IMPDH decreases the guanine nucleotide pool, which likely results in the decreased efficiency of MPXV DNA and/or RNA synthesis. In addition to this mode of action, the inhibition of the nucleic acid synthetic pathway induced the expression of interferon-stimulated genes to inhibit hepatitis C and E virus replication in Huh7 cells (17, 30, 31). We addressed this possibility by treating Huh7 cells with MPA at anti-MPXV effective concentration ranges; however, we did not observe significant induction of the representative interferon-stimulated genes ISG15 and ISG56 (Fig. S2). Consistent with the essential role of nucleic acid synthesis in the replication of most or all the virus species, MPA has been reported to inhibit a wide range of viruses, including dengue, Zika, SARS-CoV-2, hepatitis C, Lassa, and Epstein-Barr viruses (15, 19, 32–34). Thus, targeting IMPDH may contribute to the development of pan-antiviral agents beyond anti-orthopoxvirus drugs. Although host cells also require guanine synthesis for survival and function, we observed a significant window for drug concentration ranges showing anti-MPXV activity without cytotoxicity, thus indicating that IMPDH represents a realistic drug target. Actually, IMPDH-targeting agents are in clinical use for the treatment of diseases, including rheumatoid arthritis, psoriasis, and nephrotic syndrome, and are promising targets for the development of new immunosuppressants and anti-cancer agents. In this study, we identified mycophenolate mofetil, AVN-944, and merimepodib as the most potent compounds against MPXV. Further antiviral analyses under more physiologically relevant conditions, such as primary cells or animal models, would demonstrate the usefulness of IMPDH inhibition in antiviral strategies.

In conclusion, our findings suggest that IMPDH can serve as a potential target for the development of anti-MPXV agents. We found that IMPDH inhibitors exerted antiviral activities against a wide range of orthopoxviruses by inhibiting viral replication. Further studies are ongoing to demonstrate the usefulness of IMPDH-targeting agents in eliminating MPXV, with the goal of improving virus-induced pathogenesis and identifying more potent antiviral agents that target IMPDH.

## Materials and methods

### Chemical compounds

The anti-vaccinia virus compound library was prepared based on a previous study (13) by selecting compounds from the Inhibitor Library (Selleck, L1100), Anti-infection Compound Library (Selleck, L3100), and Immunology/Inflammation Compound Library (Selleck, L4100). A list of compounds in the library is presented in Table S1. Brincidofovir and AVN-944 were purchased from Cayman Chemical Company and MedChemExpress, respectively. Guanosine and ribavirin were purchased from Sigma–Aldrich. The compounds were dissolved in dimethyl sulfoxide (DMSO).

### Cell culture

An African green monkey kidney-derived cell line (VeroE6 cells) and a human hepatoma cell line (Huh7 cells) were maintained in Dulbecco’s modified Eagle’s medium (DMEM; Fujifilm Wako), which was supplemented with penicillin and streptomycin sulfate (Thermo Fisher Scientific) and 5% fetal bovine serum (FBS; Nichirei) for VeroE6 or 10% FBS for Huh7. The cells were then incubated under 5% CO2 at 37 °C.

### Compound screening

VeroE6 cells were seeded at 2 × 10^4^ cells/well in a 96-well plate. At 16 h after seeding, the cells were treated with MPXV Zr-599 (Congo Basin strain) (35) at an MOI of 0.1 and with 10 μM of each compound for 72 h. We confirmed robust cytopathology upon MPXV infection using DMSO as a control (Fig. 1A-b). We screened for compounds that protected cells from MPXV-induced cell death. Cells fixed with 4% paraformaldehyde and then stained with DAPI were counted with an ImageXpress Micro Confocal high-content imaging analyzer (MOLECULAR DEVICES), as previously described (14) (Fig. S1). Compounds that increased the number of viable cells by more than 50-fold compared to the DMSO-treated control were selected as the first hits (Fig. S1).

### Preparation of viruses

MPXV strains Zr-599 (Congo Basin strain), Liberia (West African strain), vaccinia virus (LC16m8), and cowpox virus (Brighton Red) were used as virus inocula (35). The viral titer was determined by plaque assay using VeroE6 cells, as previously described (36). Virus stocks were stored at −80 °C until use.

### Cytotoxicity assay

The cell viability assay was performed using the Cell Counting Kit-8 (DOJINDO) according to the manufacturer’s protocol.

### Indirect immunofluorescence analysis

The cells were washed with phosphate-buffered saline, fixed with 4% paraformaldehyde for 30 min, and permeabilized with 0.005% digitonin for 15 min at room temperature. Rabbit anti-vaccinia virus antibody (Abcam) and anti-rabbit Alexa Fluor Plus 555 (Thermo Fisher Scientific) were used as the primary and secondary antibodies, respectively. Nuclei were visualized using 4,6-diamidino-2-phenylindole (DAPI), and fluorescence was visualized using a fluorescence microscope (BZ-X710; Keyence). Quantification of the red fluorescence area was performed using a BZ-X Analyzer (Keyence).

### Real-time PCR/RT-PCR analysis

DNA was extracted from the cells using a QIAamp DNA Mini Kit (QIAGEN) according to the manufacturer’s protocol. Real-time PCR detection of the ATI gene of MPXV was performed using TaqMan Gene Expression Master Mix (Thermo Fisher Scientific) following the manufacturer’s instructions. The primers and probe used were as follows: forward primer, GAGATTAGCAGACTCCAA; reverse primer, GATTCAATTTCCAGTTTGTAC; and TaqMan probe, FAM-CTAGATTGTAATCTCTGTAGCATTTCCACGGC-TAMRA (35).

RNA was extracted from the cells using an RNeasy Mini Kit (QIAGEN) according to the manufacturer’s protocol. Real-time RT-PCR analysis was performed using Fast Virus 1-Step Master Mix (Thermo Fisher Scientific) following the manufacturer’s instructions. The primer and probe sets were purchased from Thermo Fisher Scientific: IMPDH1: Hs04190080_gH, IMPDH2: Hs00168418_m1, and beta actin: Hs01060665_g1.

### Time-of-drug-addition assay

VeroE6 cells were infected with MPXV at an MOI of 0.1 for 1 h in the presence (a, b) or absence (c) of the compound. After the virus inoculum was removed and washing with PBS was performed, the cells were incubated with medium supplemented with (a, b) or without (c) the compound. 1 hour later, the medium in (b) and (c) was removed, washing was performed again, and medium without (b) or with (c) the compound was added. After a further 22 h of incubation, the cells were collected to detect viral DNA by real-time PCR.

### Electron microscopy analysis

VeroE6 cells were trypsinized and fixed with buffer [2.5% glutaraldehyde, 2% PFA, and 0.1 M phosphate buffer (pH7.4)] at 4°C, followed by post-fixation with 1% osmium tetroxide, staining with 0.5% uranyl acetate, dehydration with a graded series of alcohols, and embedding with epoxy resin (37). Ultrathin sections were stained with uranyl acetate and lead citrate and observed under a transmission electron microscope. At least 150 cells per sample were observed in ultrathin sections, and representative images are shown in Fig. 5.

### RNA interference

siRNA targeting human IMPDH1, Silence Select Pre-Designed siRNA (s7413), and human IMPDH2; Silencer Validated siRNA (106308) were purchased from Thermo Fisher Scientific. An ON-TARGETplus Non-targeting Pool (D-001810-10), which was used as a negative control, was purchased from Dharmacon. Huh7 cells were transfected with 10 nM siRNA using Lipofectamine RNAiMAX according to the manufacturer’s protocol (Thermo Fisher Scientific).

### Western blot analysis

Cells were lysed with Passive Lysis Buffer (Promega), separated by SDS-PAGE with Bolt Bis-Tris Plus Gel (4-12%, Thermo Fisher Scientific), and transferred to polyvinylidene difluoride membranes using an iBlot2 instrument (Thermo Fisher Scientific). Anti-IMPDH1 rabbit polyclonal antibody (Invitrogen), anti-IMPDH2 polyclonal antibody (Proteintech), and anti-beta actin monoclonal antibody (Cell Signaling Technology) were used as primary antibodies. SuperSignal West Dura Extended Duration Substrate (Thermo Fisher Scientific) was used to visualize the signals, which were then detected with a ChemiDoc XRS instrument (Bio-Rad).

### Exogenous guanosine supplementation analysis

Huh7 cells infected with MPXV for 1 h were treated with or without 12.5, 25, or 50 μM guanosine (Sigma-Aldrich) in the presence or absence of 5 μM MPA. After 24 h of infection, the cells were recovered to detect viral DNA using real-time PCR.

### Statistical analysis

Data are presented as the mean ± standard deviation (SD). All statistical analyses were performed using Student’s *t* test. Values of \**P* < 0.05 and \*\**P* < 0.01 were considered statistically significant, and N.S. indicates not significant.

## Acknowledgments

The Huh7 cell line was kindly provided by Dr. Francis V. Chisari of the Scripps Research Institute. This work was supported by The Agency for Medical Research and Development (AMED) (JP21fk0108589, JP21fk0108421, JP22fk0310504, JP22jm0210068, JP22wm0325007), the Japan Society for the Promotion of Science KAKENHI (JP20H03499, JP61H02449), the JST MIRAI program (JPMJMI22G1), and the Takeda Science Foundation.

## Author Contributions

T.H., T.M., D.A., H.O., E.S.P., M.K., J.M., K.S., K.T., S.O., A.H.A., S.N., K.K., and K.W. performed the experiments. T.Y., M.S., K.M., T.S., H.E., and Y.T. contributed to the materials. T. H. and K. W. prepared the manuscript. T.H. and K.W. acquired funding. K.W. supervised the study.

