## Supplementary material for "Identification of inosine monophosphate dehydrogenase as a potential target for anti-monkeypox virus agents": Fig. S1, S2, Table S1

[1]Research Center for Drug and Vaccine Development, National Institute of Infectious Diseases, Tokyo 162-8640, Japan, [2]Department of Veterinary Science, National Institute of Infectious Diseases, Tokyo 162-8640, Japan, [3]Department of Pathology, National Institute of Infectious Diseases, Tokyo 162-8640, Japan, [4]Department of Virology II, National Institute of Infectious Diseases, Tokyo 162-8640, Japan, [5]Department of Applied Biological Science, Tokyo University of Science, Noda 278-8510, Japan, [6]Department of Virology I, National Institute of Infectious Diseases, Tokyo 162-8640, Japan, [7]MIRAI, Japan Science and Technology Agency (JST), Saitama 332-0012, Japan

[†¶]These authors contributed equally to this work.

[#]Corresponding author: Koichi Watashi, Ph.D.
Research Center for Drug and Vaccine Development, National Institute of Infectious Diseases, 1-23-1 Toyama, Shinjuku-ku, Tokyo 162-8640, Japan

**Table of contents**

**Fig. S1, S2**
**Table S1**

**Fig. S1**

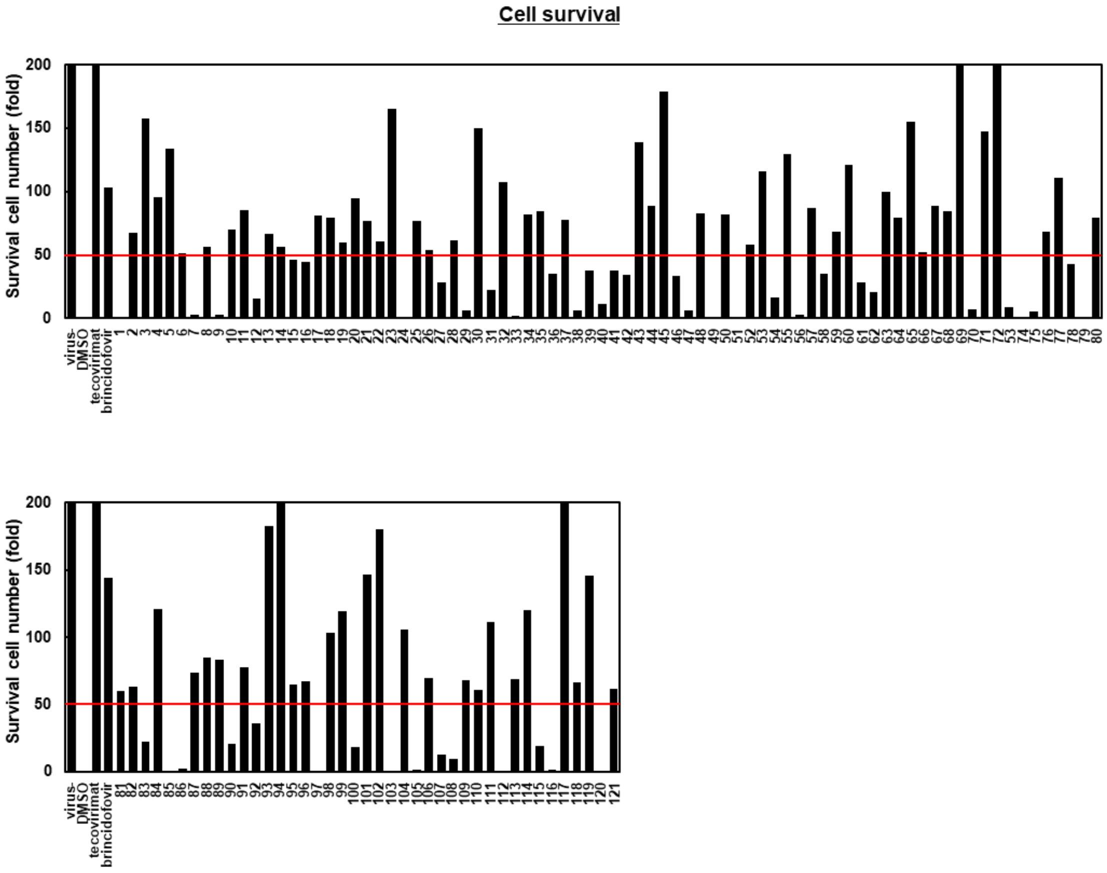


**Fig. S1. Primary screening of the compound library in MPXV-infected VeroE6 cells.** VeroE6 cells were infected with or without MPXV at an MOI of 0.1 and treated with compounds at 10 μM (2 compounds treated at 2 μM are shown in Table S1) or 0.1% DMSO. Tecovirimat and brincidofovir were used as positive controls. After 72 h of infection, the number of surviving cells was quantified by DAPI staining using a high-content imaging analyzer. The red line shows a 50-fold higher cell survival rate relative to that of the DMSO control in MPXV-infected cells.

**Fig. S2**

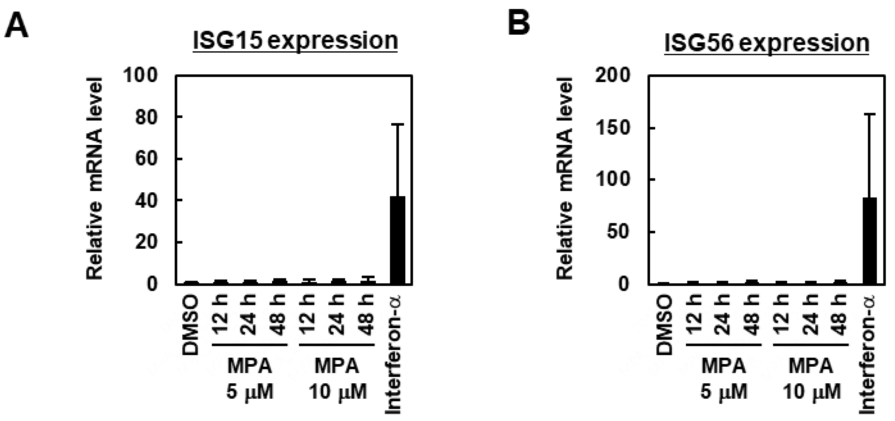


**Fig. S2. Expression of interferon-stimulated genes (ISGs) upon MPA treatment.** Huh7 cells were incubated with MPA (5 or 10 μM), interferon-α (1000 U/ml), or DMSO (0.1%). After incubation with MPA for 12, 24, and 48 h or with interferon-α or DMSO for 48 h, intracellular RNA was extracted and mRNA expression levels of ISG15 (A) and ISG56 (B) were measured by real-time RT-PCR. The Y-axis shows the value relative to that of DMSO-treated cells as a control.

**Table. S1. List of drugs in the library**

| | |
|---|---|
| 1 | ABT-737 |
| 2 | Linifanib |
| 3 | Dovitinib |
| 4 | Dasatinib |
| 5 | Gefitinib |
| 6 | Luminespib |
| 7 | MLN8054 |
| 8 | Cabozantinib |
| 9 | Mocetinostat |
| 10 | BMS-754807 |
| 11 | Tanespimycin |
| 12 | Delanzomib |
| 13 | Ganetespib |
| 14 | Onalespib |
| 15 | ABT-751 |
| 16 | BIIB021 |
| 17 | WZ8040 |
| 18 | ENMD-2076 |
| 19 | Cladribine |
| 20 | Methotrexate |
| 21 | Clofarabine |
| 22 | YM201636 |
| 23 | OSI-930 |
| 24 | Etoposide |
| 25 | KU-0063794 |
| 26 | Vincristine sulfate |
| 27 | BX-912 |
| 28 | Floxuridine |
| 29 | Genistein |
| 30 | SP600125 |
| 31 | HMN-214 |
| 32 | Fludarabine |
| 33 | Selisistat |
| 34 | Gemcitabine |

| | |
|---|---|
| 35 | Adefovir Dipivoxil |
| 36 | Azacitidine |
| 37 | Cyclocytidine HCl |
| 38 | Atorvastatin Calcium |
| 39 | Gandotinib |
| 40 | Ixazomib |
| 41 | Ixazomib Citrate |
| 42 | Avasimibe |
| 43 | OSI-420 |
| 44 | UK 383367 |
| 45 | Apigenin |
| 46 | Phloretin |
| 47 | Tolbutamide |
| 48 | Mycophenolic acid |
| 49 | MG-132 |
| 50 | OSI-027 |
| 51 | URB597 |
| 52 | PF-04929113 |
| 53 | WYE-125132 |
| 54 | ICG-001 |
| 55 | Ibrutinib |
| 56 | KW-2478 |
| 57 | Mardepodect |
| 58 | KX2-391 |
| 59 | AMG-900 |
| 60 | MK-2461 |
| 61 | Nocodazole |
| 62 | RITA |
| 63 | Vistusertib |
| 64 | Lonafarnib |
| 65 | AZD4547 |
| 66 | TAE226 |
| 67 | TPCA-1 |
| 68 | StemRegenin 1 |
| 69 | Golvatinib |

| | |
|---|---|
| 70 | ML130 |
| 71 | WHI-P154 |
| 72 | CCG 50014 |
| 73 | Niclosamide |
| 74 | Anagrelide HCl |
| 75 | Fexofenadine HCl |
| 76 | Cabozantinib malate |
| 77 | Nifuroxazide |
| 78 | PD168393 |
| 79 | Oprozomib |
| 80 | PP1 |
| 81 | XL888 |
| 82 | SC144 |
| 83 | KPT-185 |
| 84 | SKI II |
| 85 | Skepinone-L |
| 86 | KPT-276 |
| 87 | CNX-774 |
| 88 | NMS-E973 |
| 89 | Rociletinib |
| 90 | TG003 |
| 91 | PTC-209 |
| 92 | Sorafenib |
| 93 | CGP 57380 |
| 94 | AR-A014418 |
| 95 | VER-49009 |
| 96 | Triapine |
| 97 | Afatinib Dimaleate |
| 98 | Tenovin-1 |
| 99 | Tyrphostin AG 1296 |
| 100 | Butein |
| 101 | Ivacaftor |
| 102 | Vidarabine |
| 103 | Teniposide |
| 104 | Cyclosporin A |

| | |
|---|---|
| 105 | Kaempferol |
| 106 | PAC-1 |
| 107 | Azaguanine-8 |
| 108 | Bergapten |
| 109 | NSC 319726 |
| 110 | PD153035 |
| 111 | Miconazole Nitrate |
| 112 | Navitoclax |
| 113 | Cytarabine |
| 114 | Erlotinib HCl (2 μM) |
| 115 | Torin 1 (2 μM) |
| 116 | Drospirenone |
| 117 | Idoxuridine |
| 118 | Ciclopirox |
| 119 | Econazole nitrate |
| 120 | Norethindrone acetate |
| 121 | Trifluridine |

Compounds were treated at 10 μM, with the exception of 114 and 115, which were treated at 2 μM.
